# Putting the “dynamic” back into dynamic functional connectivity

**DOI:** 10.1101/181313

**Authors:** Stewart Heitmann, Michael Breakspear

## Abstract

The study of fluctuations in time-resolved functional connectivity is a topic of substantial current interest. As the term “*dynamic* functional connectivity” implies, such fluctuations are believed to arise from dynamics in the neuronal systems generating these signals. While considerable activity currently attends to methodological and statistical issues regarding dynamic functional connectivity, less attention has been paid toward its candidate causes. Here, we review candidate scenarios for dynamic (functional) connectivity that arise in dynamical systems with two or more subsystems; generalized synchronization, itinerancy (a form of metastability), and multistability. Each of these scenarios arise under different configurations of local dynamics and inter-system coupling: We show how they generate time series data with nonlinear and/or non-stationary multivariate statistics. The key issue is that time series generated by coupled nonlinear systems contain a richer temporal structure than matched multivariate (linear) stochastic processes. In turn, this temporal structure yields many of the phenomena proposed as important to large-scale communication and computation in the brain, such as phase-amplitude coupling, complexity and flexibility. The code for simulating these dynamics is available in a freeware software platform, the “Brain Dynamics Toolbox”.

## Introduction

The brain is a dynamic machine *par excellence*, tuned through the principles of self-organization to anticipate the statistics and movement of the external milieu^1,2^. Its unceasing dynamics and cycle of prediction-action-perception mark it as distinct from even the most advanced deep learning platforms despite impressive advances in machine learning. Systems neuroscience is likewise incorporating dynamic algorithms into its core methodologies^3,4^, in the design of hierarchical models of perception and inference^5^; dynamic approaches to clinical disorders^6^; dynamical models of functional neuroimaging data^7,8^ and dynamic frameworks for the analysis of resting state fMRI data^9^. Dynamic models are at the heart of the distinction between functional connectivity and effective connectivity (see Box 1)^10^ and can help disambiguate correlated activity due to mutual interactions from that caused by input from a common source.

Research into the dynamics of resting state fMRI data is currently very active, and takes its form largely through the study of non-stationarities in time-resolved functional connectivity^11-13^. However, the topic remains hotly disputed, with both positive^12,14,15^ and negative^16^ reports. In addition, fundamental statistical issues continue to be contested, including the utility of sliding-window analyses^17-19^ as well as core definitions of stationarity^20^. Another issue of substance pertains to the *causes* of putative non-stationarities (assuming they exist); in particular, whether non-stationarities reflect subtle cognitive process (random episodic spontaneous thought,i.e. “rest”^21^); whether they are slower processes that nonetheless retain cognitive salience (such as drifts in attention and arousal^22^); or whether they are nuisance physiological and head motion covariates that have been inadequately removed from fMRI time series^16^. Regardless of these debates, the over-arching motivation of the field is that resting-state brain activity is endowed with functionally relevant complex neuronal dynamics – either as the substrate for ongoing “thought”, or to prime the cortex for perception and action^23^. So, the central question seems not whether such neuronal dynamics exist, but to what extent they can be detected in functional neuroimaging data.

Dynamic models of large-scale brain activity can play a key role in this field by proposing the types of instabilities and dynamics that may be present^24-28^. The purpose of the present paper is to employ simple dynamic models to illustrate the basic processes (“primitives”) that can arise in neuronal ensembles and that might, under the right conditions, cause true nonlinearities and non-stationarities in empirical data. In doing so, we also aim to disambiguate some key terms in the field. First, the differences between non-stationarity and nonlinearity: Both can herald underlying dynamics, can cause rejection of common non-parametric nulls, and (as we will see) can occur individually or together. Second, the distinctions between key terms in dynamic systems theory, especially the catch-phrase terms of meta-stability and multistability (which are often used interchangeably). Hopefully this is a constructive step toward a more definitive resolution of the uncertainties in the field.

#### Box 1: Definitions

*Functional connectivity:* The statistical correlation between remote neurophysiological recordings (fMRI voxels, EEG channels, reconstructed sources). May be symmetrical (e.g. Pearson’s correlation coefficient) or asymmetrical (e.g. partial correlation coefficient).

*Effective connectivity:* The inferred influence of one neuronal population on another. Effective connectivity cannot be directly estimated from linear or nonlinear metrics of time series interdependence but rests upon estimation of a generative model of causal interactions^10^.

*Autonomous nonlinear system:* A time dependent dynamical system with constant (time invariant) parameters.

*Stationarity:* We adopt the simple notion of so-called *weak-sense stationarity*^20^, namely that the time-lagged auto-correlation and cross-correlation functions are invariant to time shifts. That is, if *X*_t_ and if *Y*_t_ are random processes for *t* = 1,2, …, then for any arbitrary integers *k*, *l, m* and *n*,

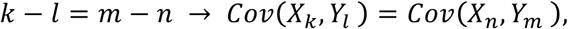

where *COV*(*X*_*i*_, *Y*_*j*_) = E[(*X*_*i*_ – *E*(*X*_*i*_)) (*Y*_*j*_ – *E*(*Y*_*j*_))] is the covariance function between *X*_*T*_ and *Y*_*T*_.

*Nonlinearity:* The behaviour of a system arises from a dynamic system with nonlinear feedback (such as a limit cycle or chaotic attractor) and cannot be closely approximated by a suitable linear reduction (as in noise-driven fluctuations near an equilibrium point).

## Methods

### 1. Coupled dynamics systems

To illustrate the breadth of synchronization dynamics, we study a system of coupled neural masses with nonlinear dynamics. This model has been previously employed to study whole brain dynamics^12,27^. The system is composed of local subsystems (or nodes) coupled together to form a larger ensemble (for review, see^3^). Each local node comprises a population of excitatory neurons and a slow variable incorporating the (simplified) response of a local inhibitory pool of neurons; Inhibitory activity is driven by local excitatory activity, to which it feeds back via a slow inhibitory current. The dynamics of neural masses are determined by a conductance-based process, with fast (instantaneous) sodium membrane currents and slower potassium currents. The dynamics within each node takes the form of a low dimensional nonlinear differential equation,

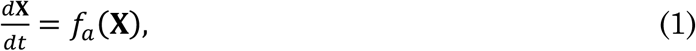

where **X** is a vector of the system’s states (cell membrane potentials, firing rates, membrane channel currents). The system has a number of physiologically-derived parameters *a*, such as synaptic connection strengths, membrane channel conductances and neural gain^29^. Depending upon the choice of these parameters, single node dynamics may range from a steady-state fixed point attractor, to fast periodic oscillations and chaotic dynamics (sustained aperiodic oscillations). Here we set local dynamics in the chaotic regime, which arise from a mixing of the time scales of the excitatory and inhibitory populations (see Appendix I).

A mesoscopic neural ensemble is constructed by permitting two or more of such local neural masses **{X**_1_, **X**_2_, … **}** to interact through a coupling function^30^. Inter-node coupling in this model is composed of excitatory-to-excitatory connectivity. These interactions are parameterized by the matrix of internode coupling **C** = **{***c*_*ij*_ **}**, where *i* is the source node and *j* is the receiver node. Connections may be reciprocal but asymmetric (*c*_*ij*_ ≠ *c*_*ji*_). Hence each node’s dynamics are governed by,

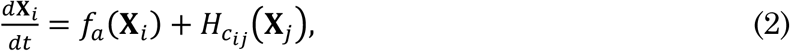

where *i* indexes the node and the coupling function *H* embodies the nature of the internode influences amongst all nodes in the system, that is the model of effective connectivity. For simulations of large empirical networks, the matrix **C** can be derived from a structural connectome such as a compilation of tracer studies or human tractography. Due to technical limitations of current state-of-the-art, connectivity matrices derived from MR-based tractography are symmetric *c*_*ij*_ = *c*_*ji*_.

Although we employ a particular model to illustrate synchronization dynamics, many of the underlying principles hold for any local system with chaotic dynamics^31^. Periodic dynamics permit a narrower range of dynamic scenarios. For most of our simulations, we focus on dyads (pairs) of coupled nodes. Complex dynamics on motifs with three or more nodes derive from the principles of two nodes, but add an additional layer of complexity, depending on their connectivity as well as the nature of axonal time delays^25,32-34^. We return to these issues below.

All simulations in this paper are performed using the Brain Dynamics Toolbox (https://bdtoolbox.blogspot.com.au/) an open-source MatLab-based toolbox for interactive simulations of neuronal dynamics (as described in Appendix II). The Brain Dynamics Toolbox allows scaling up to simulate large ensembles, the employment of other local node dynamics, the introduction of local stochastic influences and the treatment of inter-node time delays. Readers may also wish to explore The Virtual Brain^35,36^, an open source Python-based toolbox specifically for simulating whole brain dynamics according to the principles explored here.

### 2. Dynamic metrics and surrogate algorithms

We employ two metrics of inter-node interactions – the traditional Pearson's correlation coefficient, and a measure of nonlinear interdependence based upon time series forecasting methods^37,38^. These are sensitive to stationary linear correlations (traditional time-averaged functional connectivity) and stationary nonlinear interdependence, respectively. The latter estimates a (normalized) prediction error based upon forward projections of each system’s dynamic trajectory: it approaches zero for highly structured, completely predictable nonlinear time series and diverges quickly toward a maximum error of one when the time series have no structure. Crucially, the measure is sensitive to nonlinearities in the time series, possessing higher values for nonlinear time series than for random time series with the same (cross and auto-correlation) linear properties. There are two versions: self-predictions are sensitive to nonlinearities within a time-series whereas cross predictions are sensitive to nonlinear interdependences between subsystems.

Estimates of dynamic, instantaneous interactions are obtained by examining the behaviour of a phase differences. The Hilbert transform is first applied to each system’s time series, allowing an estimate of the instantaneous phase (and amplitude) of a signal^39^. The Hilbert transform of a time series *x*(*t*) is given by,

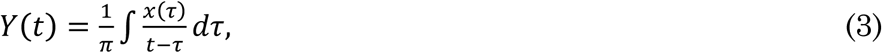

that can be used to compose the analytic signal,

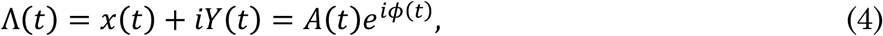

which uniquely defines the instantaneous amplitude *A*(*t*) and phase *Φ* (*t*) of the signal *x*(*t*). Phase dynamics between two signals *x*_*i*_(*t*) and *x*_*j*_(*t*) are then given by,

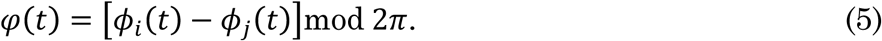

In finite length, auto-correlated time series, measures of (linear and nonlinear) sample correlations are generally not zero, even for uncoupled, independent systems. Measures of correlation taken from large numbers of samples do centre at zero, but the variance across individual samples can be substantial. To perform statistical inference on the typically modest number of data available, it is thus necessary to compare empirical measures of coupling to a null distribution derived from ensembles of surrogate data: These are pseudo-time series derived from empirical data by resampling methods that preserve the time series length, auto-correlation structure and amplitude distribution but have had the property of interest (nonstationarity, nonlinearity) destroyed. If the empirical measure falls outside of the null distribution then the data can be inferred to contain that property of interest.

For the present study, we employ a non-parametric phase-randomization method^40^. Briefly, multivariate data are mapped into the frequency domain by application of the Fourier transform. The phase of each frequency is then independently rotated by a random increment between 0 and 2π. The data are then transformed back to the time domain. By leaving the amplitude of each frequency untouched, this process preserves the power spectrum of the time series and hence the linear auto-correlations. By rotating the phases of different time series (in a multivariate stream) by the same random increment, the cross-correlations are also preserved^41^. An additional step restores the amplitude distribution of the original time series, which are otherwise rendered Gaussian^42^. This resampling approach can be adapted for complex three-dimensional data enclosed within a bounded spatial domain, such as whole brain fMRI, by using the wavelet transform^43^.

Phase randomization works because smooth dynamic trajectories generated in a low dimensional phase space (such as by equation 1) generate time series with highly structured phase relationships across frequencies. To establish the presence of nonlinearities and/or non-stationarities, we thus perform the following pipeline of randomization. To test for significant linear cross-correlations, we simply rotate the time series relative to one another (thus preserving auto- but destroying cross-correlations) and test the original against the correlations from the time rotated surrogate data. To test for nonlinearities within a single time series, we perform phase randomization and compare the nonlinear self-prediction errors of the original time series to the ensuing surrogate distribution. Finally, to establish nonlinear interdependence, we apply a multivariate phase randomization and compare the nonlinear cross predictions of original and surrogate ensemble.

## Results

We first explore the emergence of dynamic synchrony between two interacting neural masses, each with three dynamical variables **X** (*t*) = **{***V*, *W*, *Z***}** exhibiting local chaotic dynamics. We plot and analyse the time series corresponding to the average membrane potential of each system, *V*_1_ and *V*_2_. In later sections, we consider the principles underlying larger ensembles and the translation of these dynamics into the setting of noisy experimental data.

### 1. Uncoupled systems

In the absence of coupling *c*_*ij*_ = *c*_*ji*_ = 0, the two coupled neural subsystems evolve independently (Figure 1A): Due to their intrinsic aperiodic dynamics, the two systems evolve in and out of phase even if their parameters are identical. Plotting the time series of one system *V*_1_ directly against the other *V*_2_ reveals the lack of any underlying synchronization structure (Figure 1B). As a result, the difference between the two systems’ phase (modulus 2*π*) unwinds (Figure 1C). It is, however, important to note that due to the auto-correlations within each time series, the linear correlation coefficient is often not close to zero for any particular finite length sample: The correlation coefficient for the time series shown in Figure 1A is 0.08. However, the distribution of the linear correlation coefficient from an ensemble of repeated realizations of the time series is centred at zero (Figure 1D). This is a reminder that anecdotal observations of non-zero correlations can easily be misinterpreted as functional connectivity in the data, where there is none.

**Figure 1.**
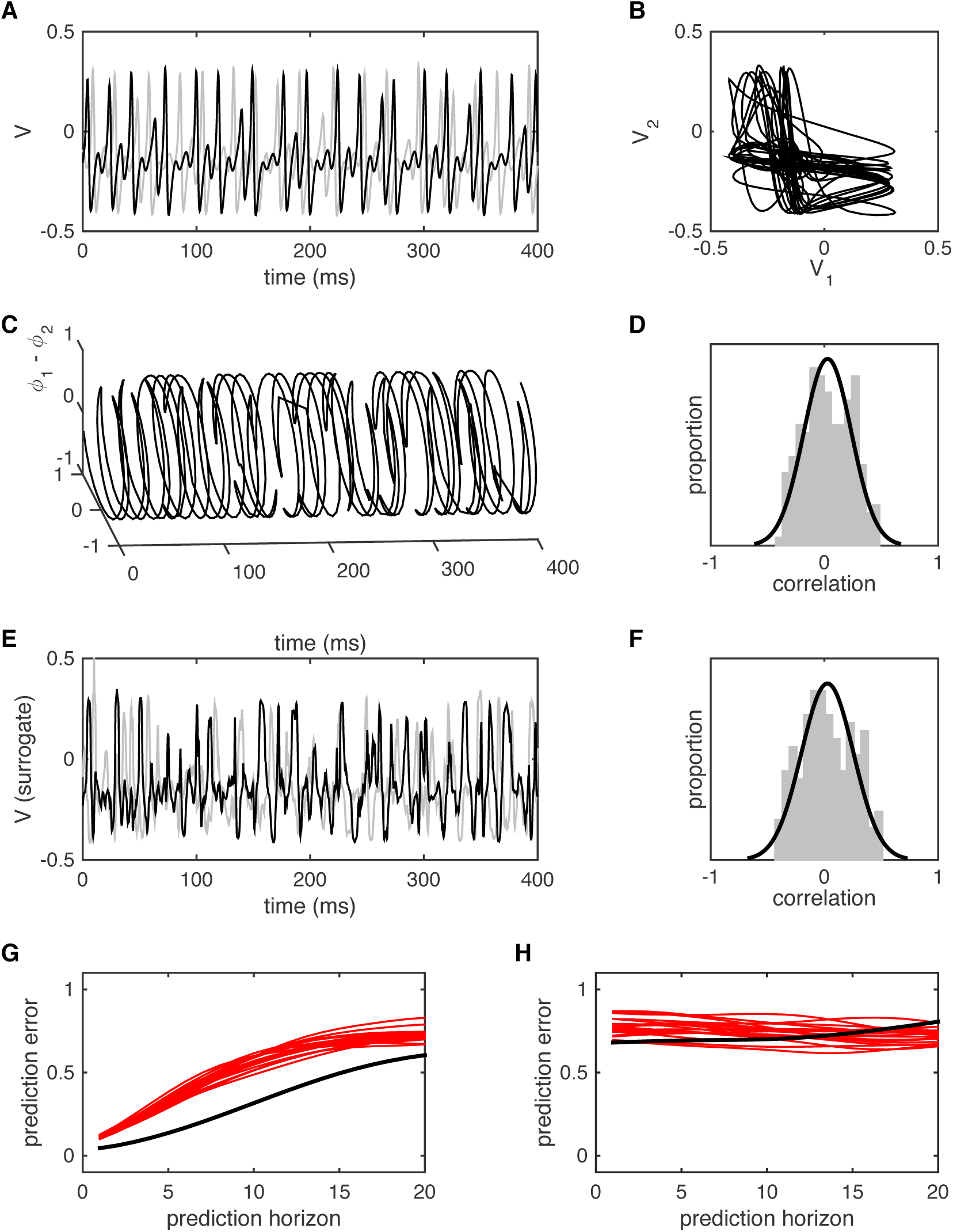
Uncoupled systems. (A) Time series for two uncoupled neural masses (V1 is black, V2 is grey) in the chaotic regime. (B) The same time series with V1 plotted against V2. Transients (t<100) have been omitted. (C) Hilbert phase of V1 relative to V2. Plotted in cylindrical coordinates with unit radius. (D) Distribution of linear correlations between V1 and V2 for multiple simulation runs with random initial conditions. (E) Amplitude-adjusted surrogates for the time series from panel A. (F) Distribution of linear correlations between surrogate data drawn from the same instances of V1 and V2 (ie one simulation run, multiple shuffles of the surrogate data). (G) Non-linear self-prediction of V1 from itself (black) and from surrogate data (red). Note that both errors grow toward one with longer prediction horizons, but the original data falls well below the null distribution. (H) Non-linear cross-prediction of V1 from V2 (black) and from surrogate data (red). Here the empirical data falls within the surrogate distribution, reflecting the absence of inter-system coupling.

Surrogate data generated from the time series in Figure 1A by (multivariate) phase randomization are shown in Figure 1E. The distribution of linear correlations between time series generated by repeated application of phase randomization are shown in Figure 1F: It can be seen that the empirical correlation (0.08) falls within the surrogate distribution. This observation confirms that the ensemble of surrogate data does adequately represent the null distribution of trivial linear correlations that arise due to the finite sample length.

Do these data contain further (i.e. nonlinear) structure? This can be tested by studying the nonlinear prediction errors, specifically how forward projections of one system’s orbits predict the actual evolution of either that same system (nonlinear self-prediction error) or the other system (nonlinear cross prediction error^37,38^). Because this approach is based upon a low-dimensional phase space reconstruction, it is sensitive to nonlinear, as well as linear correlations within the data. Here we see that such forward predictions (of one system predicting itself, Figure 1G, and of one system predicting the other, Figure 1H) are less than their theoretical maximal value of one (black lines). The nonlinear (self-) prediction errors fall well below the forward predictions arising from surrogate data (red lines), because the original time series have internal nonlinear structure, arising from the local chaotic dynamics. However, the nonlinear cross prediction errors fall within the null distribution, because there is no coupling and thus no nonlinear interdependence.

In sum, uncoupled chaotic neuronal dynamics give rise to auto-correlated time series with trivial linear cross-correlations that distribute around zero. Nonlinear self-prediction errors lie outside the null distribution, confirming that each time series contains nonlinear (chaotic) structure. However nonlinear cross-prediction errors fall within the null distribution generated by surrogate data that contain the same linear correlations. *That is, these data arise from independent (uncoupled) stationary nonlinear processes.*

### 2. Generalised synchronization

In the presence of strong unidirectional coupling, e.g. *c*_12_ = 0.6, *c*_21_ = 0, two neural subsystems with identical parameters exhibit a rapid convergence to complete synchrony: That is, the second (slave) system rapidly adjusts its dynamics to match those of the first (master) system (Figure 2A). Thereafter the two systems pursue identical orbits – that is, they exhibit *identical synchronization*, evidenced by their rapid convergence to perfect phase synchrony (Figure 2C) and their states approach the hyper-diagonal in phase space, *V*_1_=*V*_2_, *W*_1_=*W*_2_, *Z*_1_=*Z*_2_. For simplicity, we plot a two-dimensional cross section through the full dimensional phase space spanned by *V*_1_ and *V*_2_ (Figure 2C). It can be seen that the initial transient (grey line) rapidly converges onto the hyperdiagonal (black line).

**Figure 2.**
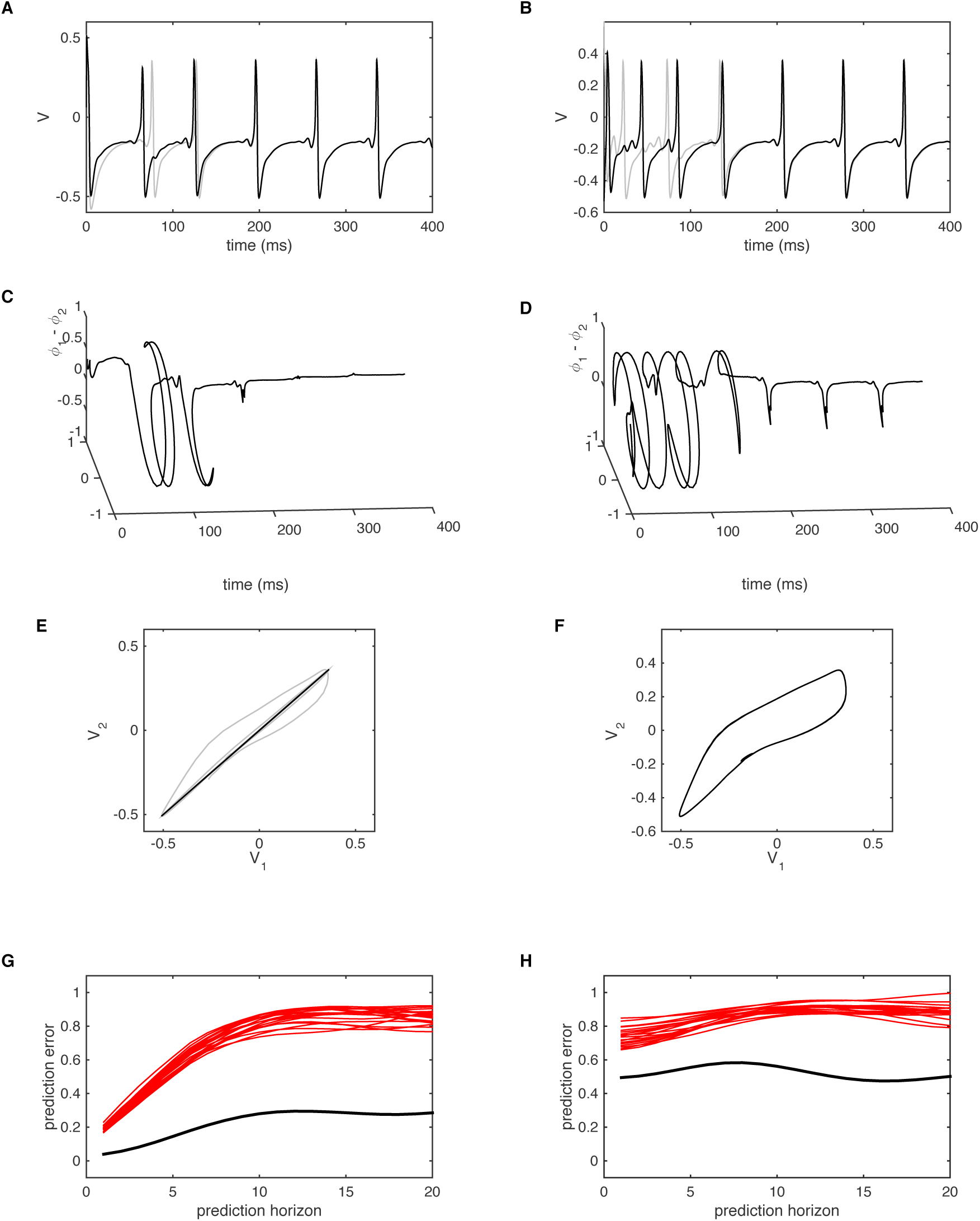
Generalised synchrony. (A) Time series for two coupled identical neural masses (V1 is black, V2 is grey) exhibiting identical synchronization. (B) Time series for two coupled non-identical neural masses (V1 is black, V2 is grey) exhibiting generalized synchronization. (C) Hilbert phase of V1 relative to V2 for the case of identical synchronization. Note the rapid approach to stable 1:1 phase synchrony. (D) Hilbert phase of V1 relative to V2 for the case of generalised synchronization. Brief, but incomplete phase slips continue to occur following the transient. (E) V1 plotted against V2 for the cases of identical synchronization. After a brief transient, the system approaches the diagonal. (F) V1 plotted against V2 for the cases of generalized synchronization. Transients have been omitted. (G) Non-linear self-prediction of V1 from itself (black) and from surrogate data (red). (H) Non-linear cross-prediction of V1 from V2 (black) and from surrogate data (red).

The onset of identical synchrony occurs for much weaker inter-node coupling if it is bidirectional, *c*_12_ = 0.05, *c*_21_ = 0.05. This is because both systems are able to simultaneously adjust their internal dynamics according to the state of the other system, leading to a more stable, integrated system.

Biological systems are obviously not composed of identical subsystems because some degree of asymmetry is inevitable. However, two neural masses with modestly mismatching parameters continue to exhibit strong, rapid and stable synchrony if the inter-node coupling is sufficiently strong (Figure 2B). These dynamics are accompanied by stable 1:1 phase locking between the two systems (Figure 2D). That is, following an initial transient of phase unwinding (until *t*=∼ 150ms), the phase difference remains close to zero, although it shows brief, bounded excursions. Rather than contracting onto the (hyper-) diagonal linear subspace, the orbits of this system converge toward a smooth manifold that lies just off the diagonal (Figure 2F). This phenomenon, known as *generalised synchronization*, arises in a broad variety of coupled asymmetric chaotic systems^44-47^. The smooth surface onto which the orbits converge is known as the *synchronization manifold.*

The time series generated in this scenario embody several instructive properties. The presence of synchrony gives rise to linear correlations that are close to unity. After a brief transient of less than 150 ms, the correlation coefficient is above 0.99 for all successive time windows. That is, the system has stationary linear cross-correlations. In the presence of static measurement noise, such a system would give rise to stationary functional connectivity (that is the ensemble linear statistics are stationary over successive time windows). However, these time series also contain deeper structure than multivariate surrogate data that possess the same linear (auto- and cross-) correlations. That is, the nonlinear prediction error (Figure 2G) and nonlinear cross prediction (Figure 2H) of the original data both lie outside of the corresponding linear null distributions. This arises because the system traverses phase space on the highly structured and smooth synchronization manifold.

In addition to the presence of stationary linear statistics, these data thus contain *nonlinear correlations* previously termed *“dynamic connectivity”*^48^. This property of the data permits rejection of the null hypothesis represented by the multivariate surrogate data, namely that the time series are generated by a stationary multivariate linear process. Since trivial analysis of the stable and very high linear correlations shows that the linear statistics are stationary, then the preceding analyses points to the (true) alternative hypothesis that the data are generated by a *stationary multivariate nonlinear process*.

### 3. Metastability

We next study the development of generalized synchrony in the presence of increasingly strong unidirectional coupling *c*_12_ > 0, *c*_21_ = 0 - that is, as the second system gradually adjusts its dynamics to those of the first. Increasing coupling *c*_12_ from zero leads to a monotonic increase in the time averaged correlation coefficient until the onset of stable generalized synchronization. However, the accompanying dynamic behaviour is quite complex^49^. When the coupling is not sufficiently strong, the two systems show instances of desynchronization, evident as a separation of the states of each system (see example in Figure 3A) and a complete unwinding of the relative phase. For weak levels of unidirectional coupling (e.g. *c*_12_ = 0.1), brief periods of generalized synchrony (and corresponding phase locking) appear amongst longer intervals of phase unwinding (Figure 3B). If the coupling is increased, the duration of synchronous epochs lengthens, and the instances of phase unwinding become confined to brief, erratic bursts (Figure 3C). Even in the presence of reasonably strong coupling (e.g. *c*_12_ = 0.5) such bursts continue to (infrequently) appear if one waits for a sufficiently long period of time (e.g. a single burst over a 20 second duration, Figure 3D). Meanwhile, as the coupling increases, the synchronization manifold contracts toward the hyper-diagonal, with asynchronous bursts corresponding to brief, disorganised, large amplitude excursions (Figure 3E).

**Figure 3.**
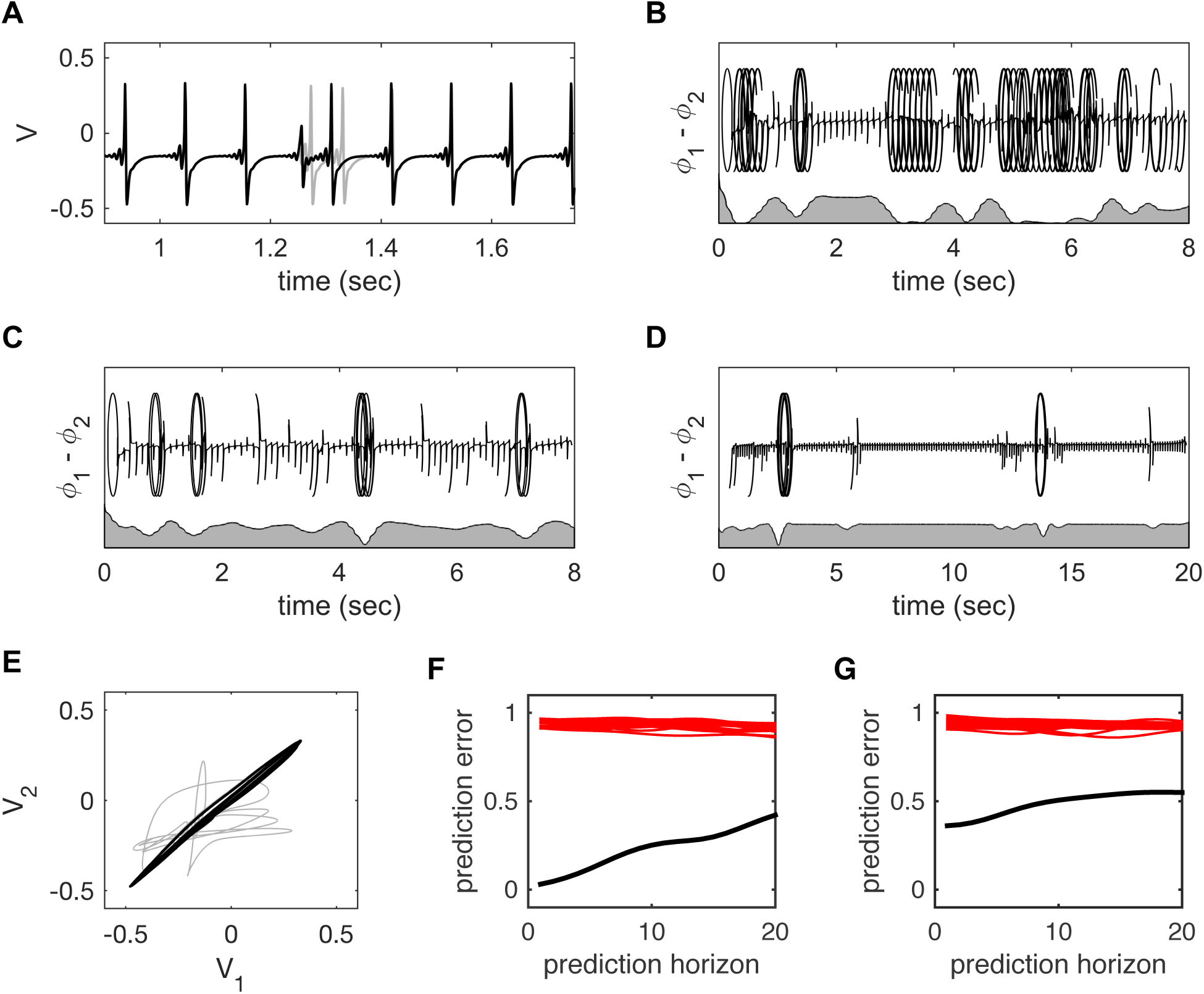
Metastability. (A) Time series for two weakly coupled neural masses (V1 is black, V2 is grey) showing a single instance of desynchronization. (B) Hilbert phase of V1 relative to V2 with relatively weak coupling. Periods of generalized synchronization are interspersed by erratic desynchronization. The grey shaded region (bottom of panel) shows the point-wise correlations between V1(t) and V2(t) smoothed over a 1 sec moving window (C) Hilbert phase of V1 relative to V2 with medium coupling. The instances of desynchronization have become relatively infrequent and briefer. (D) With strong coupling, instances desynchronization are relatively rare. (E) Plot of V1 versus V2 for the case of strong coupling. The desynchronization is seen as a brief, erratic excursion from the synchronization manifold. (F) Non-linear self-predictions of V1 from itself (black), and (G) Non-linear cross-predictions of V1 from V2 (black). Predictions of V1 from surrogate versions of V2 are shown in red. The time series retain nonlinear structure despite the instances of desynchronization.

The occurrence of such bursts corresponds to a dynamical phenomenon known as *metastability*. In brief, for strong coupling, the system possesses a single, low dimensional chaotic attractor that is embedded within the synchronization manifold: Although the dynamics of this chaotic attractor are reasonably complex (Appendix I), both systems converge onto the same manifold, corresponding to stable (and stationary) generalized synchronization^50^. The dynamics considered within the full (6 dimensional) space spanned by both systems becomes relatively simple. However, if the coupling is slowly weakened from this scenario, there appears a critical value *c*_*k*_ below which instabilities appear within the synchronization manifold and the system “blows-out” into the full phase space for brief instances (this is formally called a *blowout bifurcation*^51,52^). In the vicinity of this blowout bifurcation *c*_12_ ≈ *c*_*k*_, the intervals between asynchronous bursts can be very long, following a heavy-tailed process^52^.

Metastability is perhaps better known when there are multiple competing states^53-55^. Such a system cycles between such states, exhibiting a broad variety of synchronous behaviours (such as a variety of cluster solutions^56^. In the present setting, there is only one such unstable state and the system hence jumps away, then returns back toward the same synchronous state. This specific type of behaviour is known in the literature as *itinerancy*^57-59^. In more technical parlance, it is an example of *homoclinic itinerancy* (“homo” referring to a single system that is both attracting and repelling).

Itinerancy endows the time series with highly non-stationary properties: The unstable bursts yield a loss of phase synchrony and a corresponding local decrease in the linear correlation coefficient, both of which return to high values during the longer periods of generalized synchronization. As a result, fluctuations in time-dependent linear correlations from the original time series are greater than those arising from multivariate (stationary) surrogate data. Nonlinear prediction errors and cross prediction errors both remain outside the null distributions (from multivariate surrogate data) even if these are obtained from long windows that contain several of the bursts (Figure 3F,G).

A final summary description of these data is therefore quite nuanced. Recall that they are generated by a coupled nonlinear dynamic system whose parameters are all constant and, in particular, do not depend upon time. These data are hence generated by an *autonomous, multivariate nonlinear process*. They yield data whose nonlinear properties (for example, phase locking) are highly dynamic. The linear properties of these dynamics are also highly non-stationary – that is, they possess *fluctuating time-resolved functional connectivity*. Moreover, because the itineracy has long-tailed (non-Poisson) statistics, these properties cannot be captured by a classic finite state Markov model and hence they can be considered to violate formal definitions of weak-sense stationarity [cite].

The term “dynamic functional connectivity” is arguably a poor term to summarise these properties and to disambiguate metastability from the stationary but nonlinear properties that arise in the setting of generalised synchronization, both of which permit rejection of the stationary, linear null: We return to this issue in the Discussion.

### 4. Multistability

We consider one further dynamical scenario that yields non-trivial, dynamics interdependence between two or more systems, namely *multistability*. In a multistable system there exist two or more stable attractors: That is, there are dynamical regimes that, in the absence of noise, trap the behaviour of a system indefinitely. Multistability then arises when there is noise ζ added dynamically to the states,

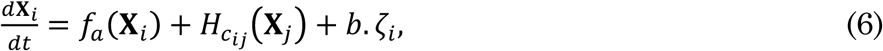

where ζ_*i*_ is a stationary zero mean stochastic process scaled in amplitude by the parameter *b*. When the noise is of sufficient amplitude, a multi-stable system is able to escape the basin of each attractor, and jump from one to the other. This is a subtle, albeit important difference between multistability and metastability: A metastable system is composed of only unstable nodes, and the evolution of the system cycles from one to the other (or back to itself) even if there is no noise ζ = 0. In contrast, a multistable will settle onto one stable attractor unless external noise is injected ζ > 0. The difference may seem subtle but the mechanisms, emergent system behaviour and resulting statistics are quite distinct (for review, see^53^).

In an array of coupled systems such as we are considering, multistability can arise when each individual node has multiple attractors. It can also emerge when the individual nodes are monostable, but the inter-node interactions introduce multiple types of synchronization dynamics^56^. In the system considered above, there is only one (chaotic) attractor per node but the coupled ensemble can exhibit multistable attractors, for example when there are three or more nodes and their interactions have axonal time delays^32^.

The emergence of multistability through the interactions of monostable elements is very interesting, but also rather complex. For reasons of relative simplicity, we will thus illustrate a system of coupled nodes where each single node has two attractors; a fixed point and a co-occurring periodic limit cycle. That is, each individual node can exhibit either steady state or oscillatory behaviour, depending on the state to which it is initially closest. A simple – or “canonical” – form of this system has been used to model the human alpha system^60^, and is a mathematical approximation to a complex neural field model^61^. The equation for the amplitude dynamics of a single node according to this simplified model are given by,

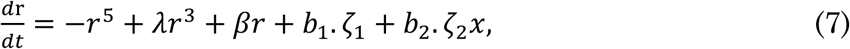

where *r* is the amplitude, *λ* and *β* are parameters that control the size and depth of the fixed point and limit cycle attractor basins. The parameters *b*_1_ and *b*_2_ control the influence of the additive ζ_1_ and multiplicative noise ζ_2_*x*, respectively (see Appendix III for full details).

When the attractor basins of each system are large (i.e. the basin boundaries are distant from the attractors) and the noise has low amplitude, the two coupled systems exhibit either noise driven low-amplitude fluctuations (Figure 4A) or high amplitude oscillations (Figure 4B). When the noise is of sufficient strength or the attractor basins are shallow, the dynamics at each node jump from one attractor to the other. In the absence of inter-node coupling, these transitions occur independently (Figure 4C). The introduction of inter-system coupling increases the coincidence in the timing of the state transitions (figure 4D). However, due to the presence of system noise, these do not always co-occur, even for relatively strong coupling.

**Figure 4.**
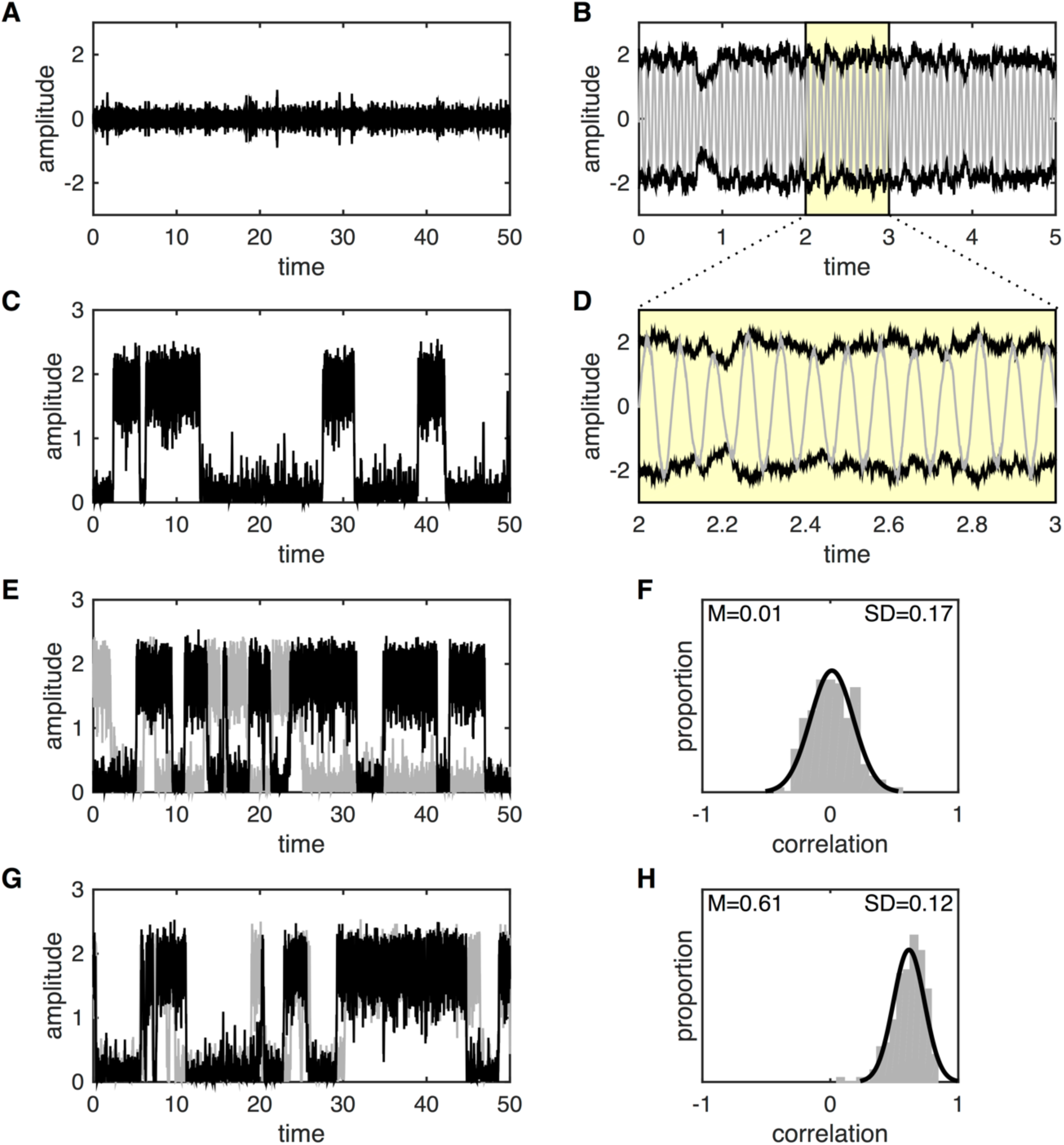
Metastability. (A) Time series of the noisy subcritical Hopf model with one node. With *β* = −10 the system exhibits a stable (noise perturbed) fixed point at *r* = 0. (B) With *β* = −6x the system exhibits a stable limit cycle with amplitude *r* = 2. Oscillations are shown in grey. Black represents the noise driven amplitude fluctuations, with close-up shown in panel D. (C) With *β* = −7.5, the system exhibits bistability with noise driven switching between the fixed point and limit cycle. For simplicity, the (grey) oscillations are not shown. (E) System with 2 nodes and *β* = −7.5 but zero coupling (*k* = 0). The systems jump between the fixed point and limit cycles independently. (F) Histogram of the linear correlations between the time-series generated by the two nodes from panel E. The simulation was repeated for N=200 trials with random initial conditions for each trial. The correlations centre at zero but with substantial inter-trial variability. (G) System with 2 nodes and *β* = −7.5 and strong coupling (*k* = 1). The jumps between the fixed point and limit cycles occur in similar time windows. (F) Histogram of the linear correlations between the time-series generated by the two nodes from panel E. The correlations centre well above zero with reduced inter-trial variability.

To illustrate the corresponding interactions between two coupled multistable nodes, we focus on their amplitude fluctuations and ignore their phases. In the absence of coupling (*c*_*ij*_ = *c*_*ji*_ = 0), linear correlations between the amplitude fluctuations converge toward zero for sufficiently long samples. However, linear correlations taken from brief samples do fluctuate considerably. Locally, the noise driven amplitude fluctuations are highly incoherent because the noisy inputs are independent (i.e. ζ_*i*_ ≠ ζ_*j*_). However, if the two systems do transition, by chance at similar times, then the local linear correlations are driven by these large amplitude changes in the variance, giving rise to large (but spurious) correlations (both positive and negative). Over time, these fluctuations centre on zero (Figure 4E), although they have high variance (SD=0.22) as a consequence of coincidental state switches. Moreover, the distribution of sample correlations (taken from short time windows) is not substantially influenced if one of the time series is randomly shifted in time compared to the other: The distribution of values is thus a reflection of the stochastic timing of the erratic amplitude jumps within each system, and whether both systems happen to switch within the same time window.

In the presence of coupling, the local fluctuations remain uncorrelated. This is due to the independence of the noise sources, ζ_*i*_ ≠ ζ_*j*_. Even though the function *f* is nonlinear, the system evolves in a largely linear fashion within each attractor, and the inter-system coupling is overwhelmed by the independent noisy perturbations around each attractor. However, if one system jumps between basins, it then exerts a strong pull on the other system, until it too jumps to the corresponding attractor. The ensuing coincidence of such large-amplitude state changes then skews the sample linear correlation toward the right (i.e. positively) so that they centre at a value greater than zero (Figure 4F). Linear correlations from long time series converge to a positive value that is typically larger than the average of the sample correlations, because such long windows are increasingly dominated by the large-amplitude states changes. Notably, the average of the sample correlations and the long-term correlation coefficient converge toward zero if one time series is independently rotated in time with respects to the other, underscoring the effect of inter-system coupling on sample correlations.

As raised above, the local (very short term) fluctuations are dominated by the independent noise sources, even in the presence of coupling. These data do not contain additional nonlinear structure (both the nonlinear prediction errors and cross prediction errors fall within the null). Between state transitions, the data resemble stationary stochastic fluctuations. Only when considered on lengthy time series data do the sample statistics reflect the presence of the underlying nonlinear multistable attractor landscape.

The time series generated by a coupled (noise-driven) multistable system hence show multivariate statistics that are *locally* stochastic, independent and stable, but are *globally* highly correlated and fluctuate substantially. If the noise term is independent of the state of the system (as per equation (6)) then the switching between attractors is Poisson^60^. The statistics of the time series can then be closely approximated by a finite state Markov process, with a fixed likelihood Ʌ of jumping states at any time, thus generating Poisson statistics with an exponential distribution of dwell times. Despite the erratic nature of the state transitions, this result thoretically renders the statistics *weak sense stationary (WSS)* because the expected correlation and cross-correlations are independent of time^20^.

However, there is one final nuance that is conceptually important. In many situations, the influence of the state noise ζ is state dependent, in which case a more general differential equation pertains,

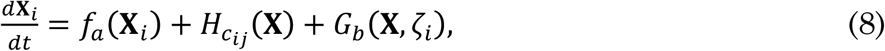

where the influence of the state noise ζ is dependent on the states **X** via the function *G*. When the noise is state dependent, (e.g. *G*_*b*_ (**X**, ζ_*i*_)= *b***X**. ζ_*i*_, as in the case of Figure 4), then the system typically gets *trapped* near each of the attractors in a non-stationary manner^60^. More technically, in setting of purely additive noise, transitions probabilities are time invariant and follow a stationary Poisson process. But with multiplicative noise, the chance of a state transition decreases as the time since the last transition increases. This non-stationarity gives rise to a heavy-tailed (stretched exponential) distribution of dwell times^60^. Long dwell times are more likely than in the case of purely additive noise case. More crucially, the dwell time is dependent on the history of the system. As a consequence, sample statistics cannot be well approximated by a standard finite state Markov process. This is a system for which the covariance between the two nodes is not time invariant and the process is thus *not* weak sense stationary.

In sum, the system governed by equation (6) for *c*_*ij*_>0 yields stochastic (linear) time series that fluctuate considerably. However, the statistics are only non-stationary in the strict sense if the noise is multiplicative (state dependent) so that systems gets trapped within each state and the ensuing statistics are non-Poisson.

### 4. Complex dynamics in larger ensembles

We have thus far restricted our analyses to coupled dyads in order to illustrate dynamic “primitives” – generalized synchronization, metastability and multistability. However, cognitive function inevitably involves exchanges between a substantial number of cortical regions – certainly more than two^62-65^. To what extent do dynamics in dyads inform our understanding of dynamics in larger ensembles, particularly as time delays (τ) between nodes become an indispensable part of modelling larger systems?

In some circumstances, the complex dynamics that occur between two nodes are inherited “upwards” when a large array of nodes are coupled together using the same principles of coupling. Thus, a system expressing multistability during the interaction between two nodes will often exhibit noise-driven multistable switching when more nodes are added. In this situation, nodes may cluster into “up” and “down” states – i.e. clusters of nodes occupying the same attractor may emerge, likewise segregated from other clusters which co-occupy a distinct state: In fact, in many coupled oscillator systems, such multistable clustering is quite generic^66^ and can theoretically encode complex perceptual information^67^.

On the other hand, introducing more nodes can lead to additional complexities and dynamic patterns that are not possible with 2 nodes. A classic example is the nature of phase relationships between nodes in the presence of time delayed coupling: With two nodes, the time delays cause a phase lag between the coupled nodes’ oscillations. However, when three nodes are coupled in an open chain (or “V”) formation, then the outer nodes can exhibit stable zero-lag synchrony, with the middle node jumping erratically between leading and lagging the outer nodes^68^. Although first described in arrays of coupled lasers^69^ considerable work has since shown that such zero-lag configurations arise in small V-shaped motifs of coupled neural systems, including spiking neurons^68^ and neural mass models^70^ Importantly, stable zero-lag synchrony between the outer nodes of a V motif can survive immersion into larger arrays, where they increase the stability of the system as a whole^28^. Such observations support the notion that these coupled triplets underlie the emergence of zero-lag correlations that have been observed in diverse neurophysiological recordings^71,72^. However, closing the 3-node motif by adding a link between the outer nodes (hence turning the V into a cycle) destroys stable zero-lag synchrony, instead promoting “frustrated” metastable dynamics^32^.

While considerable progress has been made in this area, the full armoury of complex dynamics in large systems of coupled neural subsystems is far from understood. For illustrative purposes, we consider a number of candidate scenarios that arise in larger arrays, specifically when simulating neural mass dynamics on a matrix of 47 cortical regions derived from CoCoMac^73^, a compilation of tracing studies from the Macaque brain that yields a sparse (22%) binary directed graph (available in the Brain Connectivity Toolbox^74^). We employ a coupling function 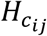 that mimics a competitive (agonist) feedback between local self-excitation and input from distant regions such that as the influence of external nodes is scaled up by a coupling constant *k*, local recurrent feedback is correspondingly scaled down by (1-*k*). All inter-node influences occur through the coupling matrix *C*_*ij*_.

The neural mass model employed here possesses time two scales – a fast local field oscillation of approximately 100Hz nested within a slower time scale of approximately 10Hz (due to the slow inhibitory feedback, see Appendix I). When parameterised with strong inter-node coupling (e.g. *k*=0.75) and a time delay that approaches the period of the fast oscillations of the neural mass model (τ = 6-10 ms), the ensemble dynamics break into a number of phase coupled clusters (Figure 5A,B).

**Figure 5.**
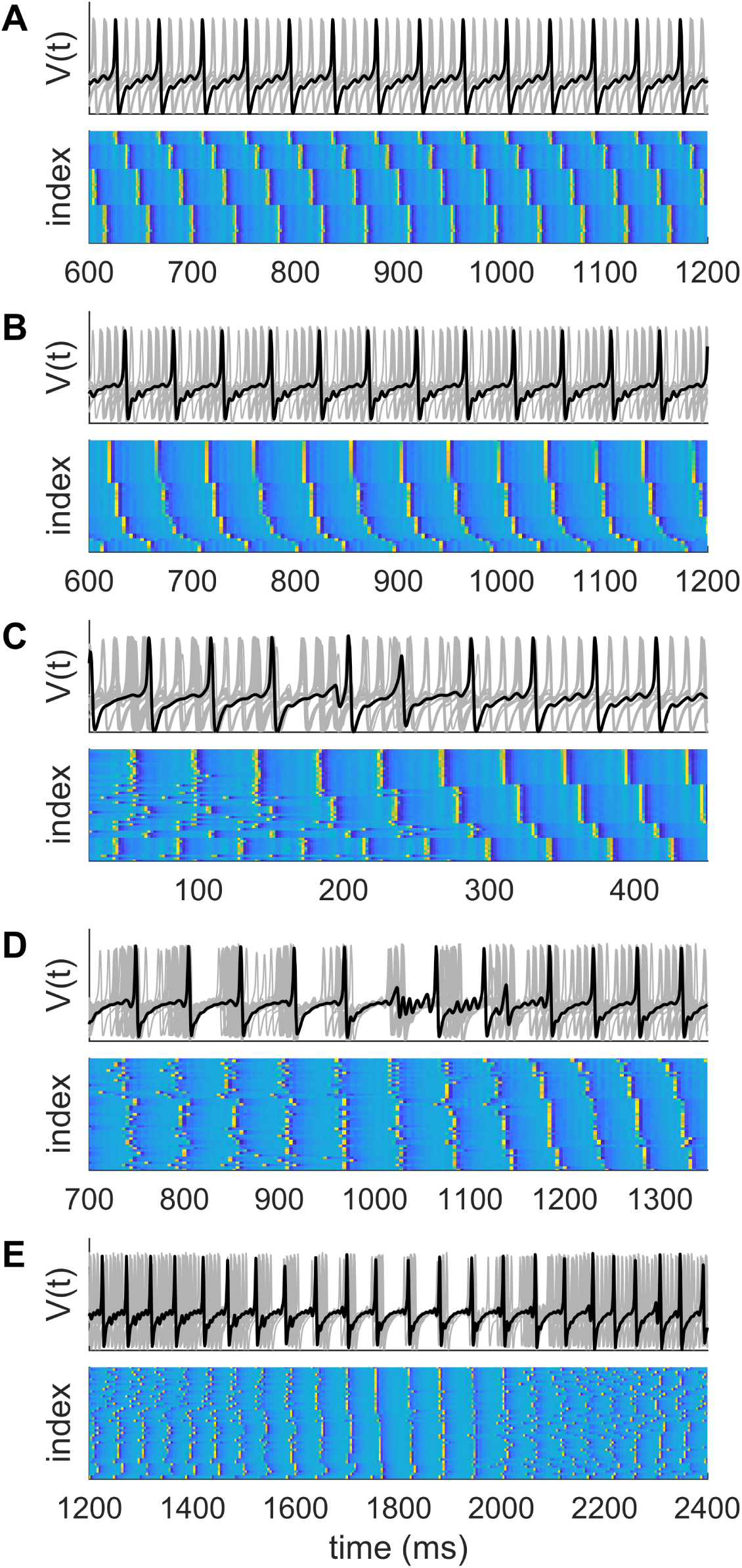
Complex dynamics in larger ensembles. (A) Stable partitioning of ensemble dynamics into 4 phase-coupled clusters with *τ* = 10ms and coupling *k*=0.75. (B) Partitioning of ensemble dynamics into 6 phase-coupled clusters with *τ* = 6ms and coupling *k*=0.75. There is slightly greater disorder in some of the clusters compared to those in panel A. (C, D) With weaker coupling and/or shorter time delays (*τ* = 5.5ms, *k*=0.ij), there are brief phase slips, leading to a reorganization of the cluster configuration. With briefer time delays (*τ* = 5ms), clustering does not occur. Instead the system shows instances of global synchrony interspersed amongst spatiotemporal chaos.

Each cluster is constituted by phase entrainment to a common beat of the faster rhythm. The full array of cluster states then recur over the course of the slow oscillation. Note that the number of clusters may differ according to the time delay (4 clusters for τ = 10ms, figure 5A and 6 clusters are apparent for τ = 6ms, figure 5B). In this scenario, the nodes within clusters show stable, stationary generalised synchronization. Nodes in different clusters also show generalised synchronization, albeit with a constant phase offset. This is an ensemble equivalent of stable generalized synchrony in coupled dyads.

This example illustrates the self-organization of coupled neural systems into dynamic communities, an example of functional segregation. Of interest, if the coupling is weaker (e.g. *k*=0.45) or the time delay shorter (τ < 6 ms), the ensemble dynamics show brief instances of desynchronization, such that the global coherence of regions into clusters decreases and nodes switch alliances between clusters (Figure 5C). Similar occasions of desynchronization can herald a reconfiguration from a poorly organised state to highly clustered dynamics (Figure 5D). In these settings, the ensemble shows some dynamic flexibility in addition to segregation. Such instances render the ensemble statistics non-stationary around the point of transition.

If a shorter time delay (τ∼ 5 ms) is incorporated into the model, then a clean demarcation into distinct phase coupled clusters does not arise. The ensemble rather shows instances of near global synchrony interspersed by longer periods of relatively dispersed dynamics (Figure 5E). During the epochs of high global synchrony, zero-lag synchrony emerges and as a result, the ensemble has highly ordered (low entropy) dynamics: Outside of these windows, the amount of order amongst the nodes is low. The ensemble statistics in this setting are both nonlinear and non-stationary.

These scenarios illustrate the capacity of neuronal ensembles to exhibit a diversity of dynamic behaviours that yield nonlinear and nonstationary statistics. In some scenarios, dynamical primitives that characterize coupled pairs (generalized synchronization, metastability and multistability) dominate the ensemble dynamics, yielding their characteristic dynamic fingerprints. New phenomena also appear, including zero lag synchrony (despite the presence of time delays) and clustering. Typically, these new behaviours compliment the basic dynamics present in coupled dyads, hence metastable bursts yielding spontaneous reconfiguration of cluster states.

## Discussion

The growth of interest in “dynamic” resting state functional connectivity motivates a deeper understanding of synchronization dynamics in neural systems. Our objective was to illustrate the breadth of synchronization dynamics that emerge in pairs and ensembles of coupled neural populations, and to propose these as putative causes of empirical observations. To recap, coupled dyads exhibit several basic forms – dynamic ‘primitives’ – that yield non-trivial statistics in the ensuing time series: Generalized synchronization yields stationary nonlinear time series; Metastable dynamics, which arise when the inter-system coupling is below a critical threshold, yield nonstationary and nonlinear statistics. Multistability yields a non-stationary process that is locally linear (i.e. on short time scales) but evidences strong nonlinear properties globally (over long time scales). When such pairs are integrated into a larger ensemble and the coupling is imbued with time delays, then these basic primitives combine with new phenomena, such as phase clustering, to cause complex dynamics that spontaneously switch between different network configurations. This yields time series whose statistics violate the assumptions of a stationary stochastic process and which hence yield non-trivial fluctuations in time resolved functional connectivity. The dynamics primitives of generalized synchronization, metastability or multistability may thus account for the spontaneous fluctuations observed in resting state fMRI data.

It is also interesting to consider the computational potential of these synchronization dynamics. Neural mass models describe the local average of neural activity, namely local field potentials and average spike rates, not individual spikes. It has been proposed that coherent oscillations in the local field potentials of disparate brain regions promotes information transfer^75^ and spike time dependent plasticity^76^. Accordingly, the dynamics illustrated in this paper would allow such neuronal binding^77^ to occur across multiple time scales, and amongst dynamic cortical assemblies that form and dissolve through active, nonlinear processes^21,78^. Dynamic synchronization, and desynchronization, could also underlie the spontaneous shifts in attention the co-occur with changes in neuronal oscillations^79,80^ and in the absence of changes in task context^22^. The nesting of a fast oscillation in a slower one – as occurs for our neural mass model - yields phase-amplitude and phase-phase interactions, which have been proposed supporting cognitive processes requiring complex spatial and temporal coordination^81-84^. More deeply, the presence of weak instabilities (such as brief desynchronizations) in cortical dynamics have been proposed as a means by which cortex can be primed to respond sensitively to sensory perturbations and thus to optimise perceptual performance^85,53^. Future work is required to elucidate the effects of exogenous stimulation and endogenous parameter modulation on neural dynamics, and thus to more directly address these issues^86^.

There are several important caveats of the present study. Most notably, functional connectivity denotes correlations between neurophysiological recordings (see Box 1): We have interrogated the time series of simulated neuronal states *X*. Neural states are not directly evident in empirical functional imaging data, which rather arise from noisy and indirect observation processes. We can somewhat crudely represent this as,

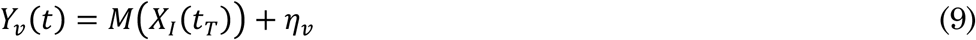

Where *Y*_*v*_ is the empirical signal in channel/voxel *v*, *M* is a complex measurement process (a nonlinear convolution over a set of regions *i* ∈ *I* and time τ ∈ *T*) and η_*v*_ is the added measurement noise. Functional connectivity is defined as *Cov*(*Y*_*v*_, *Y*_*w*_) not *Cov*(*X*_*i*_, *X*_*j*_.) In the case of fMRI, the BOLD signal arises from the summed effect of neuronal activity signalling slow changes in blood flow, mediated by neurovascular coupling^87^. The nett effect of this hemodynamic response function (HRF) is a broad low pass filter^88^, which also includes some spatiotemporal mixing if sampled at sufficiently high spatial resolution^89^. Empirical functional connectivity in rs-fMRI experiments thus reflect slow changes (<0.1Hz) in synchronization dynamics plus the effect of local spatial mixing. Although we do not explicitly model the observation function, it is worth noting that both meta- and multistability yield fluctuations that are substantially slower than the time scales of the single node neural dynamics (see Fig 3B-D), clearly extending into the slow time scales of the HRF. Explicitly incorporating an observation function into a computational framework is crucial to any definitive resolution of fMRI fluctuations. However, the appearance of slow fluctuations in synchrony from fast dynamics lies at the core of the body of work using neural mass models to study fMRI^12,27,28,90-92^.

In comparison, EEG and MEG directly detect field fluctuations and do not suffer the same temporal filtering as fMRI. Analyses of resting state EEG^93-95^ and MEG^96^ data using nonlinear time series techniques has shown that the human alpha rhythm is imbued with substantial nonlinear structure. Integrating these findings with use of biophysical models has shown that the alpha rhythm arises from multistable switching between a fixed point and periodic attractor in the presence of multiplicative noise^60,61^ – precisely the scenario illustrated in Figure 3. This process yields fluctuations in alpha power whose time scales clearly extend into those of the HRF^97^. Crucially, the presence of multiplicative noise in the model (and non-Poisson dwell times for each attractor) imply, as discussed above, that the system statistics are history dependent and are not (weak sense) stationary according to formal, quite restrictive definitions^20^. By this we can infer that the statistics of resting state cortex *are* non-stationary.

Despite their superior temporal fidelity, EEG and MEG sensor data involve substantial spatial summation of source activity. Although “off the shelf”^98,99^ source reconstruction methods are now available, they inevitably incorporate assumptions about the covariance structure of the sources and the measurement noise^100^. As such, and despite post-hoc unmixing steps (orthogonalization^101^), there does not yet exist a definitive account of the contribution of synchronization dynamics to source-level M/EEG activity. Given recent accounts of complex multi-network switching in such data^102^, substantial steps toward this end seem within reach.

In addition to spatial and temporal filtering, empirical data are also corrupted by extraneous “noise”, including physiological effects (EMG, respiratory confounds) and measurement noise (thermal fluctuations etc). These external (nuisance) noise sources (η in Equation 9) are conceptually distinct from intrinsic system noise (ζ in Equation 7), which are an essential part of neuronal dynamics. While resolving the contribution of measurement noise to resting state fluctuations is largely a methodological and empirical issue^16,103^, computational modelling can also assist. As we have seen above, multistable and metastable processes yield specific heavy-tailed statistics. Most nuisance confounds either have a specific time scale (such as respiratory effects^104^), or have very short-range correlations (such as thermal effects). The hope here is to use the statistical fingerprints of synchronization dynamics to help disambiguate true from spurious fluctuations in observed data. Given the salience of brain-body interactions (as indexed by physiological and behavoural interdependences^105^), it should also be considered that *some* physiological correlates will index true neuronal dynamics^106^ and not simply artifacts. Computational models that incorporate somatic and physiological dynamics – which thus embody and not merely eschew these signals – may be required here^107^.

Complex network dynamics are a topic of substantial interest^108^, particularly in computational and network neuroscience ^109-112^. Network fluctuations co-occur with a variety of fluctuating cognitive processes within and across rs-fMRI sessions ^113,114^ To understand how basic dynamics between pairs of coupled systems scale up, we simulated networks dynamics on a structural connectome using connectivity data from Cocomac. This bottom-up approach^28,32^ contrasts with typical to-down approaches in the field – namely of embracing emergent network dynamics without recourse to the dynamics amongst the basic elements ^12,27,90-92^. Our simulations showed how dynamical primitives mix with new ensemble phenomena to inform global dynamics, including the presence of clustering and global synchronization. We did not explore the specific role of the CoCoMac connectome network topology in shaping these dynamics, nor correlations between functional and structural connectivity, which has been the subject of substantial prior work^115^. However, modelling work in this field – mirroring empirical resting state research – has focussed on structural correlates of time-averaged functional connectivity. Future work is required to build upon the early forays in this direction^28,92^.

Although it has intuitive appeal, the term “dynamic functional connectivity” is arguably a clumsy one as it suggests, perhaps naively, that dynamic processes exist within the stream of observable data. Dynamics occur in the underlying neuronal system. If they extend into the temporal and spatial aperture of a particular functional neuroimaging modality (in sufficient strength to survive corruption by measurement effects) then they cause non-trivial statistics in time resolved data samples. From this perspective, it would be preferable to use simple descriptive terms to capture the fluctuating properties of these time-resolved data and reserve the notion of dynamics to refer to their underlying causes.

## Acknowledgements

This manuscript was supported by the National Health and Medical Research Council (118153, 10371296, 1095227) and the Australian Research Council (CE140100007).

## Appendix I: Neural mass model

Full details of this model can be found in Ref^29^. The version here is adapted from Ref^6^.

This simple neural mass model has three state variables: mean membrane potential of local pyramidal cells *V*, mean membrane potential of inhibitory interneurons *Z*, and the average number of open potassium ion channels *W*. Conceptually, as a conductance-based neural mass model, its formulation is similar to the Morris-Lecar single neuron above, but where the quantities are now population means, and with the addition of feedback from a passively enslaved local inhibitory population. The governing equations are,

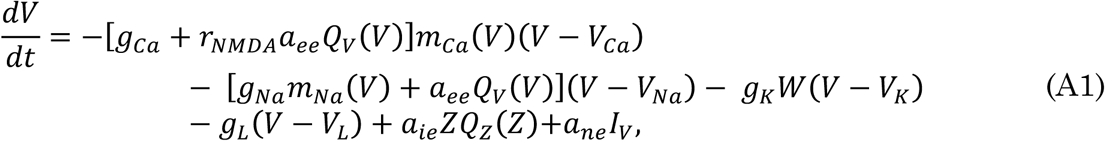

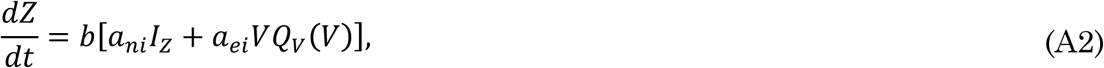

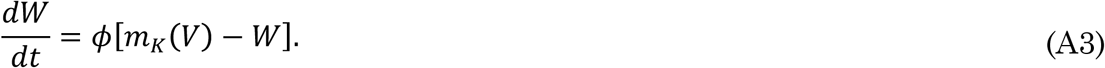

These are in nondimensional units where capacitance *C*=1; time is thus also nondimensional but usually considered to be numerically equivalent to milliseconds^29^. Here *I*_*V*_ and *I*_*Z*_ are nonspecific inputs to excitatory and inhibitory populations, respectively; in the absence of noise we set *I*_*V*_ = *I*_*Z*_ = *I*_*0*_. The model distinguishes between AMPA and NMDA channels, where *r*_NMDA_ denotes the ratio of NMDA receptors to AMPA receptors, and *a*_*xy*_ terms parameterize the strength synaptic coupling from population *x* (= *e*, *i*, *n*, where *n* is a non-specific input) to population *y* (= *e*, *i*) (note that this index ordering is the reverse of the physics convention). Parameters *b* and *Φ* are rate parameters (inverse time constants) that determine the time scales of *Z* and *W*, respectively. Setting b=0.1 means that the time scale of the inhibitory cells, *Z* are slow with respects to the pyramidal cells, *V.*

As in the Morris-Lecar neuron, for the pyramidal neurons the *g*_*ion*_ terms are conductances of the corresponding ion channels, and the *m*_*ion*_(*V*) functions describe the voltage-dependent gating of these ion channels, except that in this mean-field case they describe population-level fractions of open channels. They take the sigmoidal form,

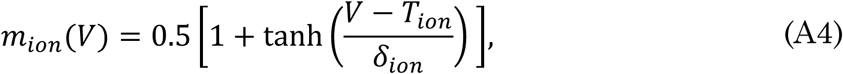

where *T*_*ion*_ is the threshold membrane potential for a given ion channel, and δ_*ion*_ is the corresponding standard deviation in this threshold. The self-feedback of the pyramidal cells is split into a conventional voltage-dependent term for sodium channels *a*_*ee*_*Q*_*V*_ (*V*) (*V* – *V*_*Na*_), and a state-dependent NMDA term, *r*_*NMDA*_ *a*_*ee*_*Q*_*V*_ (*V*) *m*_*Ca*_ (*V*) (*V* – *V*_*Ca*_) This more complex term incorporates the state-dependent nature of NMDA-gated calcium channels.

The voltage-dependent functions *Q*_*V*_ and *Q*_*Z*_ are the mean firing rates of the excitatory and inhibitory populations, respectively, also given by sigmoidal forms

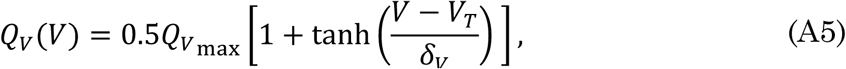

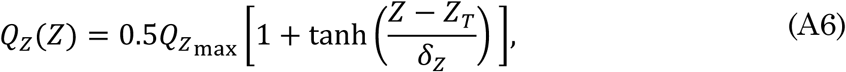

where 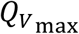 and 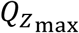 are the maximum firing rates of the excitatory and inhibitory populations, respectively, and *V*_*T*_ and *Z*_*T*_ are the corresponding thresholds for axon potential generation, and *δ*_*V*_ and *δ*_*Z*_ are the standard deviations in these thresholds. Figure A2 shows a representative plot of *Q*_*V*_(*V*).

Note that the inhibitory population *Z* does not possess conductance-based membrane dynamics, but rather is passively slaved to the pyramidal population: its membrane potential (and resulting firing rate) is driven by the firing rate of the output of the pyramidal cells, responding with the slow time scale parameterized by the factor *b*. The inhibitory population thus acts as a passive low-pass filter of the pyramidal cells. Even in the absence of a leaky current, the inhibitory neurons thus do not saturate, but rather oscillate with the slow time scale of the system (approximately 1/10^th^ of the fast pyramidal cells for the parameters used here). It is this mixing of time scales that leads to the chaos evident in Figure 1.

Parameters are given in Table A1. To introduce additive noisy input to the excitatory population, we set *I*_*V*_ = *I*_*0*_ + *ση*(*t*), where *η* is zero-mean Gaussian white noise and unit variance, and *σ* is the standard deviation of this component. To introduce multiplicative noisy input to the excitatory population, we set *I*_*V*_ = *I*_*0*_ + *σV*(*t*) *η*(*t*).

**Figure A1:**
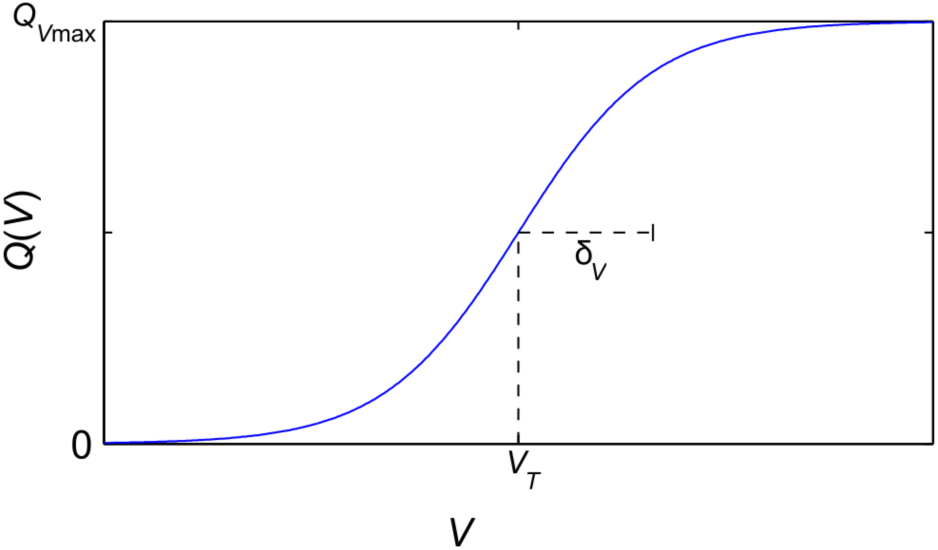
Sigmoidal voltage-dependent firing-rate function *Q*_*V*_(*V*) with threshold potential *V*_*T*_ and threshold standard deviation *δ*_*V*_.

**Table A1:** Parameters for neural mass model (in nondimensional units)

| Parameter | Description | Value |
| --- | --- | --- |
| $g_{Ca}$ | Maximal conductance of calcium ion channel | 1.1 |
| $g_{Na}$ | Maximal conductance of sodium ion channel | 6.7 |
| $g_K$ | Maximal conductance of potassium ion channel | 2 |
| $g_L$ | Conductance of passive leaky membrane | 0.5 |
| $V_{Ca}$ | Equilibrium potential of calcium ion channel | 1 |
| $V_{Na}$ | Equilibrium potential of sodium ion channel | 0.53 |
| $V_K$ | Equilibrium potential of potassium ion channel | -0.7 |
| $V_L$ | Equilibrium potential of passive leaky membrane | -0.5 |
| $T_{Ca}$ | Threshold for calcium ion channels | -0.01 |
| $T_{Na}$ | Threshold for sodium ion channels | 0.3 |
| $T_K$ | Threshold for potassium ion channels | 0 |
| $\delta_{Ca}$ | Threshold standard deviation for calcium ion channels | 0.15 |
| $\delta_{Na}$ | Threshold standard deviation for sodium ion channels | 0.15 |
| $\delta_K$ | Threshold standard deviation for potassium ion channels | 0.3 |
| $V_T$ | Firing threshold potential for excitatory population | 0 |
| $Z_T$ | Firing threshold potential for inhibitory population | 0 |
| $\delta_V$ | Firing threshold standard deviation for excitatory population | 0.7 |
| $\delta_Z$ | Firing threshold standard deviation for inhibitory population | 0.7 |
| $Q_{V_{\max}}$ | Maximum firing rate for excitatory population | 1 |
| $Q_{Z_{\max}}$ | Maximum firing rate for inhibitory population | 1 |
| $b$ | Rate constant for inhibitory dynamics | 0.1 |
| $\phi$ | Rate constant for potassium dynamics | 0.7 |
| $r_{NMDA}$ | Ratio of NMDA to AMPA receptors | 0.25 |
| $a_{ee}$ | Synaptic strength from excitatory to excitatory | 0.36 |
| $a_{ie}$ | Synaptic strength from inhibitory to excitatory | 2 |
| $a_{ne}$ | Synaptic strength from nonspecific to excitatory | 1 |
| $a_{ei}$ | Synaptic strength from excitatory to inhibitory | 2 |
| $a_{ni}$ | Synaptic strength from nonspecific to inhibitory | 0.4 |
| $I_0$ | Mean nonspecific input strength | 0.3 |

## Appendix II: The Brain Dynamics Toolbox.

The Brain Dynamics Toolbox (https://bdtoolbox.blogspot.com.au/) is an open-source toolbox for simulating and exploring problems in dynamical systems using Matlab. It includes a graphical user interface with which users can explore the behaviour of a dynamical system in real-time by manipulating system parameters. It supports the most common classes of differential equations used in computational neuroscience; namely Ordinary Differential Equations, Delay Differential Equations and Stochastic Differential Equations.

The figures in the present paper were generated using version 2017c of the Brain Dynamics Toolbox. It can be downloaded from http://bdtoolbox.blogspot.com.

The time-series in Figure 1 were generated using the BTF2003 model which simulates a network of recurrently connected neural masses as described by Breakspear, Terry and Friston^29^. The complete source-code for the model is shipped with the toolbox and can be run from the matlab command-line as follows:

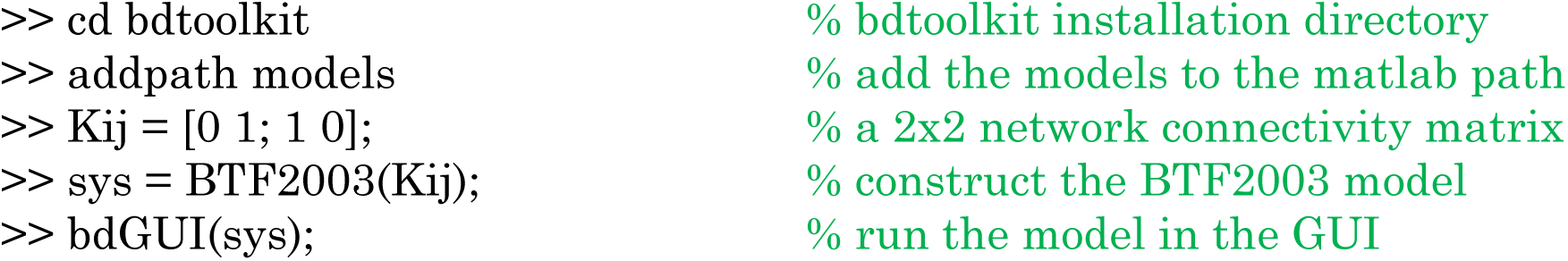

The parameters and initial conditions of the model can be manipulated using the graphical controls on the right-hand side of the application window. For figure 1, all parameter values are the default with the exception of (*I* = 0.2; *δ*_*Z*_ = 0.8, *g*_*Ca*_ = 0.9) which put the system in a “faster” regime (and hence more clearly illustrate subsystem independence). These are reset by changing the corresponding parameter values and rerunning the simulation

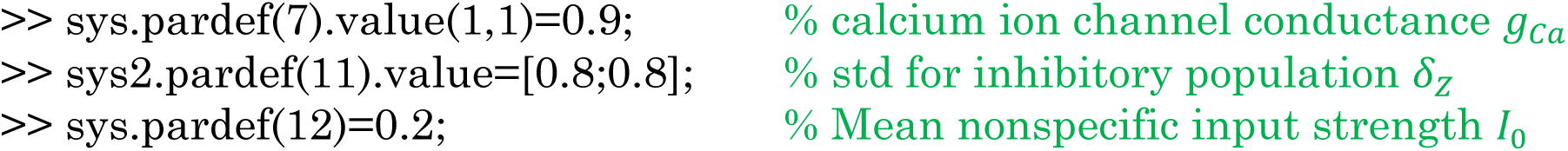

Panel A is replicated by the Time-Portrait panel of the graphical user interface. Panel C is replicated by the Hilbert Transform panel. Panel E is replicated by the Surrogate data panel available from the pull-down menus (New Panel - Surrogate Transform).

The time-series in Figure 2 were likewise generated using the BTF2003 model but in this case, the coupling has been “turned on” (C=0.1) and the default settings (Appendix I, Table 1) are used for all parameters. The metric for computing the non-linear prediction errors (panels F and G) are not included in the toolbox.

The simulations in Figures 4A-D were conducted using the FRRB2012 model which reproduces the canonical model of multistability proposed by Freyer, Roberts, Ritter and Breakspear^60^. It too is shipped with the toolbox and can be run as follows:

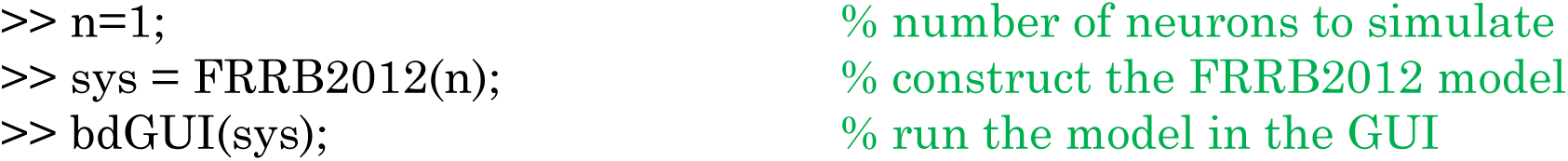

The parameter values for Figure 4A were rho=0.65, eta=1, beta=-3 and lambda=4. Figure 4B used the same values except that beta=-1. Likewise for Figure 4C which had beta=-2. Figures 4E,G were simulated using the FRRB2012b model which is a network variant of the FRRB2012 model that is also shipped with the toolbox. It can be replicated as follows:

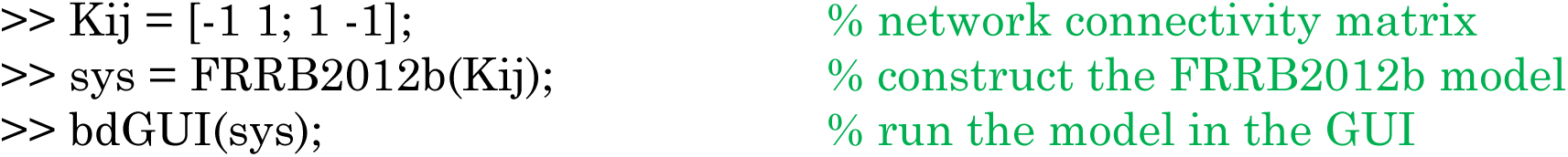

Where the global coupling coefficient was k=0 and k=0.5 in Figures 4E and 4G respectively. All other parameters were lambda=4, beta=-2, eta=1, rho=0.65.

The simulations in Figure 5 were conducted using the BTF2003DDE model which is a time-delayed variant of the BTF2003 model^29^. In this case it uses the 47x47 macaque connectivity matrix from the Brain Connectivity Toolbox which can be loaded as follows:

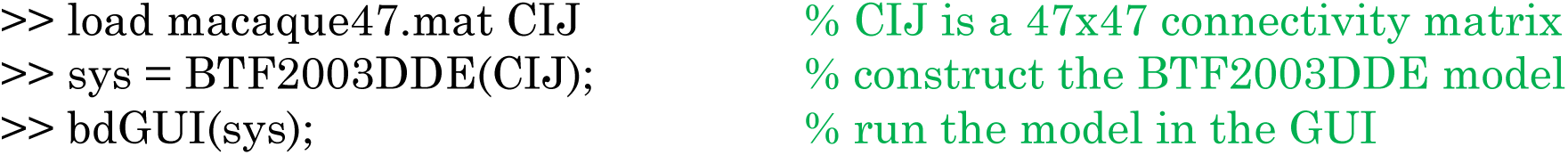

The coupling strength and time delay can be manipulated with the options in GUI.

## Appendix III: Bistable Hopf model

Full details of this model can be found in Ref^60^. The version here is adapted from Ref^6^.

This model is a type of *normal form* model that describes a generic limit cycle oscillator with bistability ^60^. Normal forms describe the dynamics near a bifurcation, and are useful because all instances of that bifurcation (whatever the system) can be reduced to essentially the same simple canonical form. Depending on the choices of its parameters, the bistable Hopf model describes a fixed point (stable or unstable) and 0, 1, or 2 limit cycles. A Hopf bifurcation is a type of bifurcation where a fixed point loses stability, giving rise to a limit cycle (also termed a periodic orbit). A model with a limit cycle is often termed an “ oscillator”. There are two types of Hopf bifurcation: supercritical, where the limit cycle is stable and coexists with the unstable fixed point, and subcritical, where the limit cycle is unstable and coexists with the stable fixed point. The Hopf normal form is most conveniently expressed in polar coordinates (*r*, *θ*), where *r* is the amplitude (so *r* = 0 corresponds to a fixed point), and *θ* is the phase. To drive a transition from the fixed point to the limit cycle (i.e., “ seizure onset”), we include independent additive *b*_1_. ζ_1_ and multiplicative *b*_2_. ζ_2_*x* noisy drives, where ζ_1_ and ζ_2_ are zero-mean unit-variance Gaussian white noise, and *b*_1_and *b*_2_ are the standard deviations of the stochastic inputs.

The state variables evolve according to,

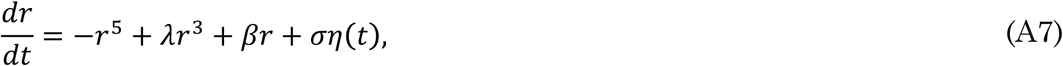

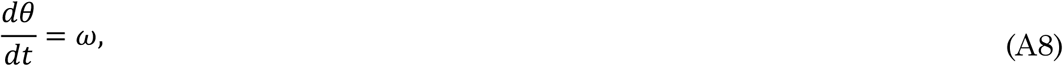

where *λ* and *β* are parameters that determine the number, amplitude, and stability of limit cycle solutions, *ω* is a constant angular velocity (in this simple case, the phase plays no role in the dynamics – it is directly proportional to time). For the bistable model here, we have extended the more typical 3rd-order Hopf bifurcation normal form to 5th order. This has the effect of introducing an additional high-amplitude limit cycle that coexists with the low amplitude one, as well as an additional bifurcation (a “ saddle-node”) where two solutions meet and annihilate. This form is bistability generic, in the sense that it emerges from a normal form that exists near any Hopf bifurcation.

